# Increased Infectivity of Human Cytomegalovirus Strain TB40/E Conferred by Variants of the Envelope Glycoprotein UL128 and the Regulatory Protein IE2

**DOI:** 10.1101/2025.04.09.647953

**Authors:** Xuan Zhou, Linjiang Yang, Giorgia Cimato, Giada Frascaroli, Ana Águeda-Pinto, Laura Hertel, Wolfram Brune

## Abstract

Human cytomegalovirus (HCMV) infects various cell types in its human host, and this broad tropism plays a vital role in viral transmission, dissemination, and pathogenesis. HCMV strains differ in their ability to infect and replicate in different cell types, but the genetic determinants of cell tropism have only begun to be understood. A widely used HCMV strain, TB40/E, contains a mixture of genetically distinct virus variants. Only a few passages in ARPE-19 epithelial cells resulted in the selective enrichment of a substrain, termed TB40/EE, which infected epithelial cells more efficiently than the parental TB40/E and induced the formation of large multinucleated syncytia. Herein, we used sequence comparison and genetic engineering of a TB40/E-derived bacterial artificial chromosome clone, TB40-BAC4, to demonstrate that the high infectivity of TB40/EE and its ability to induce syncytia in epithelial cells depends on two single-nucleotide variants (SNVs) affecting the envelope glycoprotein UL128 and the major viral transactivator protein, IE2. While the intronic SNV in UL128 increased splicing of the *UL128* transcript, it surprisingly decreased viral infectivity and replication in epithelial cells. The additional introduction of the IE2 SNV reversed this phenotype, increasing infectivity and syncytium formation. This SNV resulted in a D390H substitution and increased the levels of several early and late viral proteins, suggesting that it altered the ability of IE2 to activate viral genes. The same two SNVs increased the ability to infect THP-1-derived macrophages and JEG-3 trophoblast cells. These results demonstrate that HCMV cell tropism depends on both envelope glycoproteins and regulatory proteins.

**Importance:** Different genetic versions of human cytomegalovirus (HCMV) affect its ability to infect various human cell types. Here we focused on a commonly used strain, TB40/E, which contains a mix of virus variants. After growing it in epithelial cells, a specific variant called TB40/EE became dominant. This variant infected epithelial cells more effectively and caused the formation of large, fused cells (syncytia). In this study, we discovered that two small genetic changes were responsible for this behavior. One change affected a protein on the viral envelope (UL128) by altering how its RNA was processed. Surprisingly, this change reduced the virus’s ability to spread, but a second change in a regulatory protein (IE2) reversed that effect. Together, these changes enhanced the virus’s ability to infect not only epithelial cells but also macrophages and placental cells. This study highlights how small genetic tweaks can influence how HCMV targets different types of human cells.

## Introduction

Human cytomegalovirus (HCMV) is a β-herpesvirus that is highly prevalent among the human population worldwide (1). HCMV infection is usually mild or asymptomatic in otherwise healthy people. However, it can cause severe disease in immunocompromised individuals, such as transplant recipients, and immunologically immature fetuses and newborns (2, 3).

HCMV exhibits a broad cell and tissue tropism in the human host, infecting various organs and tissues (4). In vitro, it replicates in diverse cell types, including fibroblasts, endothelial, epithelial, and smooth muscle cells (5). Placental trophoblast cells were also shown to be susceptible to HCMV infection (6–8), suggesting that the infected trophoblast may be involved in HCMV transmission from mother to fetus. Among hematopoietic cells, HCMV shows a specific tropism for myeloid lineage cells (5). Myeloid progenitor cells and monocytes serve as a reservoir for latent virus as they are non-permissive for lytic replication (9). However, infected monocytes can circulate in the bloodstream and migrate into tissues, thereby contributing to virus dissemination in the host. Monocyte differentiation into macrophages or dendritic cells may lead to virus reactivation from latency, replication, and infection of neighboring cells (10–12).

HCMV cell tropism is regulated by viral envelope glycoproteins mediating receptor binding and entry (13). For viral entry, the viral envelope needs to fuse with the host cell membrane. This process is driven by the core fusion machinery consisting of glycoprotein B (gB), the primary fusogen, and gH/gL complexes that regulate gB activity (14). Viral entry into fibroblasts is mediated by the trimeric gH/gL/gO complex, which interacts with platelet-derived growth factor receptor alpha (PDGFRα) on the cell surface (15, 16). Infection of endothelial cells, epithelial cells, and macrophages requires binding of the pentameric gH/gL/UL128/UL130/UL131 complex to cellular receptors such as Neuropilin-2 or OR14I1 (17, 18). The relative amount of trimeric and pentameric complexes on the viral envelope is highly variable and depends on the specific HCMV strain as well as on the cell type from which the viruses emerge, thus broadening the virus’s ability to infect specific tissues (19). Not surprisingly, the HCMV fusion machinery also plays a crucial role in cell-cell fusion and syncytium formation, which is induced to varying degrees by different HCMV strains (20).

Once HCMV DNA is delivered to the nucleus of permissive cells, lytic DNA replication can occur. It depends on six core proteins necessary for DNA synthesis: the viral DNA polymerase and its accessory protein (encoded by *UL54* and *UL44*, respectively), the helicase-primase complex (*UL105, UL70, UL102*), and the single-stranded DNA-binding protein (*UL57*) (21). These proteins accumulate in membraneless nuclear compartments, which are formed by liquid-liquid phase separation and provide optimal conditions for viral gene expression and replication (22). In addition, the 86-kDa immediate-early 2 (IE2) protein and the UL84 protein are required for viral DNA replication (21, 23). IE2 is the major viral transactivator protein, and it is also involved in viral DNA replication (24). The UL84 protein associates with the HCMV lytic origin of DNA replication (*ori*Lyt) in infected cells, probably by interacting with IE2 or the UL44 protein (25–28). While UL84 is essential for replication in most HCMV strains, it is dispensable for others such as TB40-BAC4 and TR (29, 30). Interestingly, UL84-independent replication is conferred by a single-nucleotide variant (SNV) in the IE2-encoding UL122 open reading frame, resulting in a H390D substitution in IE2 (30).

HCMV propagation in cell culture results in mutations ranging from single-nucleotide substitutions to larger alterations such as deletions, duplications, and rearrangements. While mutations are thought to occur in a stochastic fashion, there is a strong selection pressure for mutations conferring a growth advantage in certain cell types or growth conditions (31–33). During passage in fibroblasts, HCMV is prone to acquiring inactivating mutations in genes *UL128*, *UL130,* and *UL131,* possibly because these genes encode components of the pentameric glycoprotein complex that are crucial for the infection of endothelial and epithelial cells but dispensable for the infection of fibroblasts (34–36). Hence, as a consequence of extensive propagation in fibroblast cultures, many laboratory strains have lost the broad cell tropism characteristic for clinical HCMV isolates. Nowadays, scientists often prefer to use genetically more authentic strains such as TB40/E, VR1814, TR, or Merlin (37–40). As the genomes of these strains have been cloned as bacterial artificial chromosomes (BACs) in *E. coli,* they can be modified easily and precisely by BAC recombineering (41, 42). TB40-BAC4, a TB40/E-derived BAC clone (43), is one of the most widely used HCMV strains. It retains endothelial and epithelial tropism even after passage in fibroblasts, presumably because it expresses relatively low levels of the pentameric complex (33, 44). While TB40-BAC4 is genetically pure, the clinical isolate TB40/E is a mixture of numerous genetic variants (39), one of which was captured in TB40-BAC4 (43). Propagation of TB40/E for only a few passages either in epithelial cells or fibroblasts has led to the isolation of two substrains, which were named TB40/EE and TB40/EF, respectively (45). TB40/EE infects ARPE-19 retinal pigment epithelial cells very efficiently and spreads rapidly in these cells. It also induces the formation of large multinucleated syncytia (45). In contrast, TB40/EF is poorly infectious for ARPE-19 cells and not syncytiogenic. Both substrains have been analyzed by deep sequencing, revealing nonsynonymous SNVs in genes UL26, UL69, UL128, UL130, US28, and a frameshift in UL141 (46). Some SNVs appeared to be enriched in TB40/EE as compared to TB40/EF, suggesting that they might be important for replication and spread in epithelial cells. However, their specific contributions to the TB40/EE-specific phenotype have remained unexplored.

In this study, we show that the combination of two TB40/EE-specific SNVs in UL122 (encoding IE2) and UL128 is responsible for the high infectivity of TB40/EE and its ability to induce cell-cell fusion in epithelial cells. The same two SNVs are also responsible for the high infectivity of TB40/EE for THP-1-derived macrophages and JEG-3 trophoblast cells. While the TB40/EE-specific SNV in UL128 improves splicing efficiency, it is not sufficient to substantially increase UL128 protein levels in viral particles. However, the IE2 H390 variant present in TB40/EE increased the expression of several early and late viral proteins, including UL128, suggesting that a single amino acid change can modify the function of the IE2 transactivator.

## Results

### Phenotypic and genetic differences between TB40/EE and TB40-BAC4

Passaging of HCMV strain TB40/E on epithelial cells and fibroblasts yielded two virus stocks named TB40/EE and TB40/EF, respectively (45). TB40/EE replicates efficiently in epithelial cells and produces large multinucleated syncytia, whereas TB40/EF does not (45, 46). To identify the genetic determinants of these TB40/EE-specific properties, we first compared TB40/EE with TB40-BAC4, the most widely used TB40/E-derived BAC clone. TB40-BAC4 is genetically pure, fully sequenced, and can be modified by BAC recombineering. Multistep replication kinetics experiments revealed that TB40/EE replicated in ARPE-19 epithelial cells faster and more efficiently than TB40, the virus reconstituted from TB40-BAC4 (Fig. 1A). TB40/EE also formed large multinucleated syncytia in ARPE-19 cells with a circular arrangement of nuclei akin to the petals of a flower (Fig. 1B). In contrast, syncytia were rare and small in TB40-infected cells. To better quantify syncytium formation, we used a previously described dual split protein (DSP) system consisting of split GFP and split *Renilla* luciferase (47, 48). ARPE-19 cells stably expressing either DSP1-7 or DSP8-11 were mixed at a 1:1 ratio, seeded, and infected with TB40/EE and TB40. Cell-cell fusion was quantified by measuring *Renilla* luciferase activity in cells at three days post-infection (dpi). As shown in Figure 1C, TB40/EE was a very strong inducer of cell-cell fusion and syncytium formation, whereas TB40 was not.

**Figure 1.**
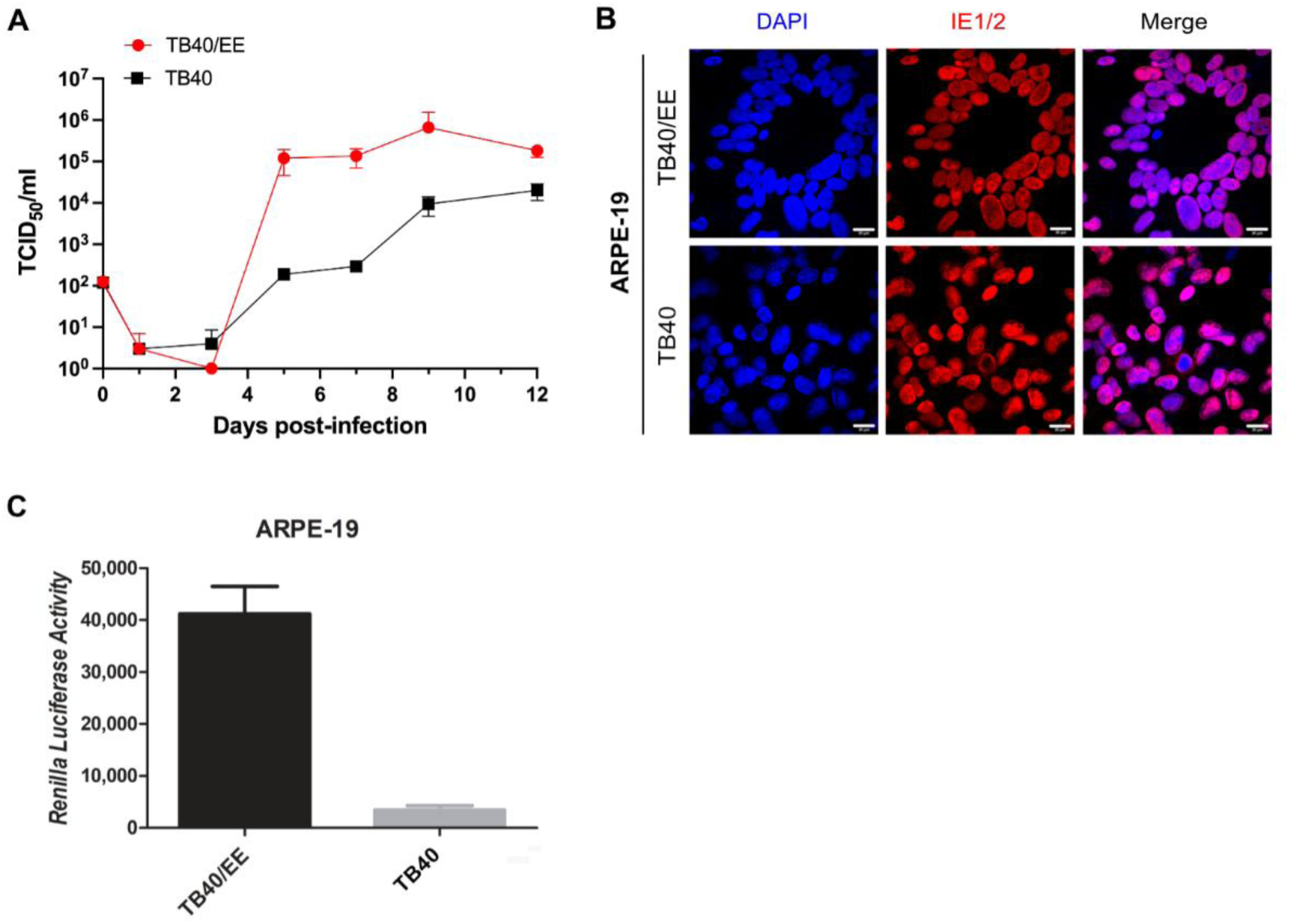
HCMV replication and syncytium formation in ARPE-19 cells. (A) ARPE-19 cells were infected with HCMV strains TB40/EE or TB40 (the TB40-BAC4-derived virus) at an MOI of 0.01. Supernatants were collected at multiple times post-infection and titrated on MRC-5 cells to determine viral titers. Data represent the mean ± SEM of three biological replicates. (B) ARPE-19 cells were infected at an MOI of 1 and fixed 5 dpi for immunofluorescence staining with an anti-IE1/2 antibody. Nuclei were counterstained with DAPI. Scale bar, 20 µm. (C) DSP-expressing ARPE-19 cells were infected at an MOI of 1. *Renilla* luciferase activity was quantified at 5 dpi. Mean ± SEM of three independent experiments.

HCMV strain TB40/E and its substrains, TB40/EE and TB40/EF, are known to be mixtures of different virus variants rather than genetically pure viruses (46). To identify the genetic differences responsible for the remarkable properties of TB40/EE, we compared the TB40/EE consensus sequence (GenBank MW439039) with the TB40-BAC4 sequence (GenBank EF999921). Apart from lacking genes US1 to US6 due to BAC vector insertion (43), the TB40-BAC4 sequence differs in only 11 positions from the TB40/EE sequence (Table 1). Of these, only two are in coding regions: an SNV in UL26 of TB40/EE leading to a lysine instead of a glutamic acid residue at position 98 (E98K) of the UL26 protein and an SNV in UL122 leading to an aspartic acid instead of a histidine at position 390 (D390H) of the IE2 protein.

**Table 1.** Nucleotide differences between TB40-BAC4 and TB40/EE.

| Region | Gene | Protein | Position* | Consequence | TB40-BAC4 | TB40/EE |
| --- | --- | --- | --- | --- | --- | --- |
| Coding region | UL26 | UL26 | 65305 | E98K | C | T |
|  | UL122 | IE2 | 203650 | D390H | C | G |
| Non-coding region | UL128 | UL128 | 209051 | / | T | G |
|  | TRL | / | 33362 | / | A | C |
|  | TRL | / | 33646 | / | G | C |
|  | TRL | / | 33754 | / | T | G |
|  | TRL | / | 33817 | / | G | T |
|  | UL21A-22A | / | 59736 | / | A | C |
|  | UL23-24 | / | 61575 | / | / | A |
|  | TRL | / | 226932-226930 | / | - | 35 nt |
|  | TRL | / | 227059 | / | G | C |

\*Coordinates refer to TB40-BAC4 (GenBank EF999921).

### SNPs in UL128 and UL122 jointly increase syncytium formation and infectivity in epithelial cells

Next, we used site-directed mutagenesis of TB40-BAC4 to determine which of the identified genetic differences might be responsible for the TB40/EE phenotype. We focused on the two SNVs affecting the coding regions of UL26 and UL122 and on the SNV in intron 1 of the envelope glycoprotein gene UL128, as this change was previously shown to affect splicing of the UL128 transcript (32). While the UL128 protein is part of the pentameric complex required for epithelial cell infection, UL26 is an inhibitor of the transcription factor NF-κB (49), and IE2 is a major transactivator protein required for early and late gene expression. Interestingly, the IE2 D390H polymorphism was previously reported to determine the requirement of the viral UL84 protein for viral replication (30).

To evaluate if these three SNPs, individually or in combination, might be responsible for the TB40/EE phenotype, we constructed seven TB40-BAC4 mutants carrying one, two, or three TB40/EE-specific SNVs affecting the NF-κB inhibitor UL26 (N), the IE2 protein (I), or the pentamer component UL128 (P). A schematic depiction of these mutants is shown in Figure 2A. The integrity of the mutant BACs was initially verified by restriction fragment length analysis and sequencing of the mutated site and subsequently by whole BAC sequencing.

**Figure 2.**
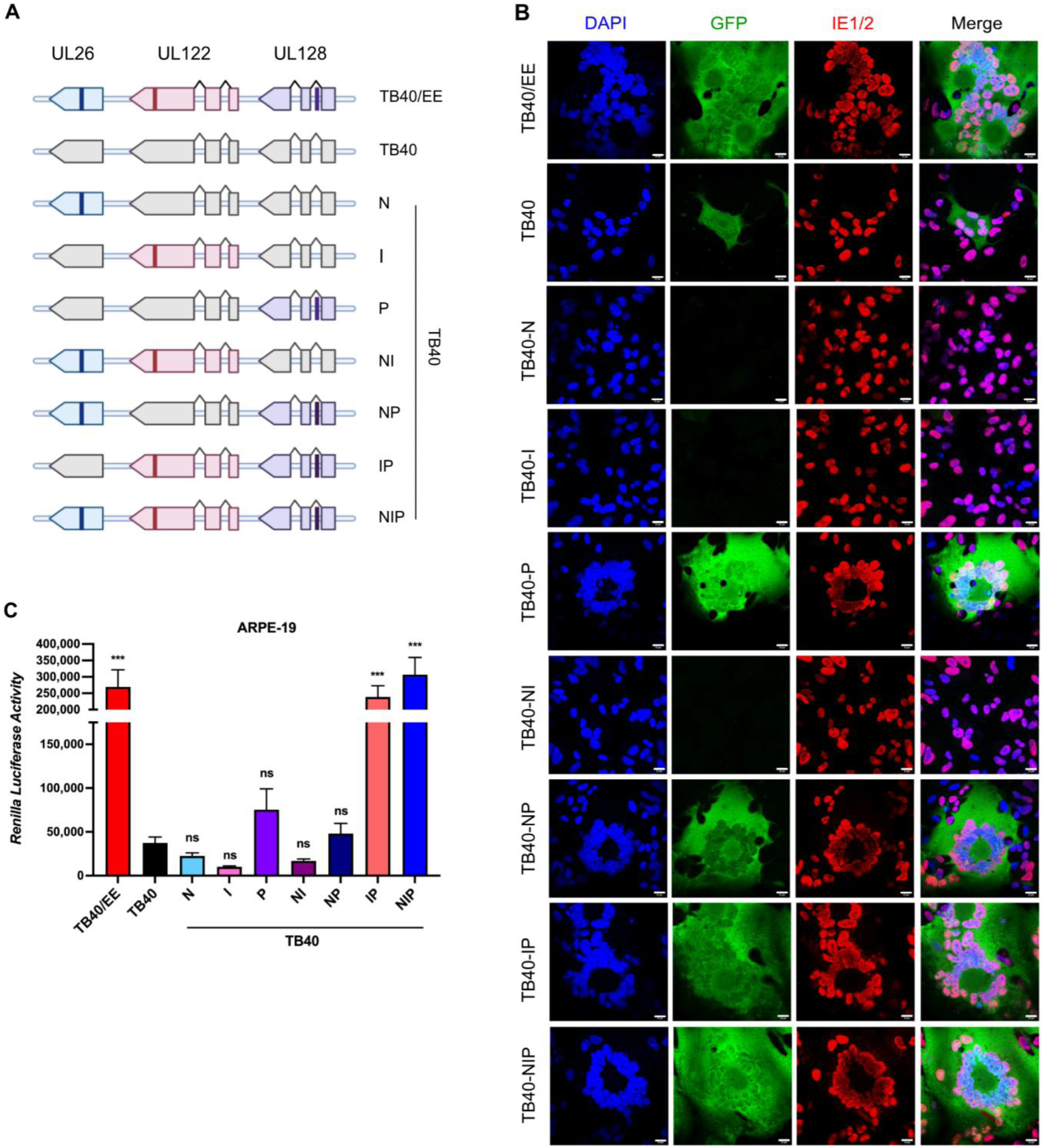
Cell-cell fusion induced by TB40/EE and recombinant TB40 strains. (A) Schematic representation of recombinant TB40 strains generated by BAC mutagenesis. TB40/EE-specific variants of the NF-κB inhibitor UL26 (N), the IE2 transactivator encoded by *UL122* (I), and the pentamer component UL128 (P) were introduced into TB40-BAC4. (B, C) DSP-expressing ARPE-19 cells were infected with TB40/EE and recombinant TB40 mutants at an MOI of 3. (B) Syncytia were visualized using the DSP system. Fused cells express the complete RLuc-GFP and are green fluorescent. IE1/2 proteins were detected by indirect immunofluorescence, and nuclei were counterstained with DAPI. Scale bar, 20 µm. (C) Cell-cell fusion was quantified by measuring *Renilla* luciferase activity at 3 dpi. Mean ± SEM of three independent experiments. Recombinant TB40 strains were compared to WT TB40. Significance was determined by one-way ANOVA with Dunnett’s multiple comparison test. ***, *P* < 0.001; ns, not significant.

First, we tested the ability of the recombinant viruses to induce cell-cell fusion and syncytium formation in ARPE-19 epithelial cells. The DSP system allowed us to image green fluorescent syncytia and quantify cell-cell fusion by measuring *Renilla* luciferase activity (Fig. 2B and C). The parental TB40 virus caused the formation of very few small syncytia (example shown in Fig. 2C). Such small syncytia were occasionally also seen in cells infected with TB40-N, TB40-I, and TB40-NI. A small number of larger syncytia were observed in ARPE-19 cells infected with the recombinant viruses TB40-P and TB40-NP. However, only TB40-IP and TB40-NIP induced the formation of numerous large syncytia similar to those generated by TB40/EE (Fig. 2B). A quantification of cell-cell fusion induced by the different viruses is shown in Figure 2C. These results indicated that SNVs in UL128 and UL122, but not UL26, are required to reproduce the strong syncytial phenotype of TB40/EE.

Next, we determined the relative infectivity of the recombinant viruses on ARPE-19 cells. Cells were infected at an MOI of 1 (based on titers determined on HFF cells) and the percentage of IE1/2-positive cells present at 2 and 4 dpi was determined by immunocytochemistry (Fig. 3). The TB40-IP and TB40-NIP mutants had the highest relative infectivity, comparable to that of TB40/EE, suggesting that the same SNVs were responsible for increased cell-cell fusion and increased infectivity in ARPE-19 cells.

**Figure 3.**
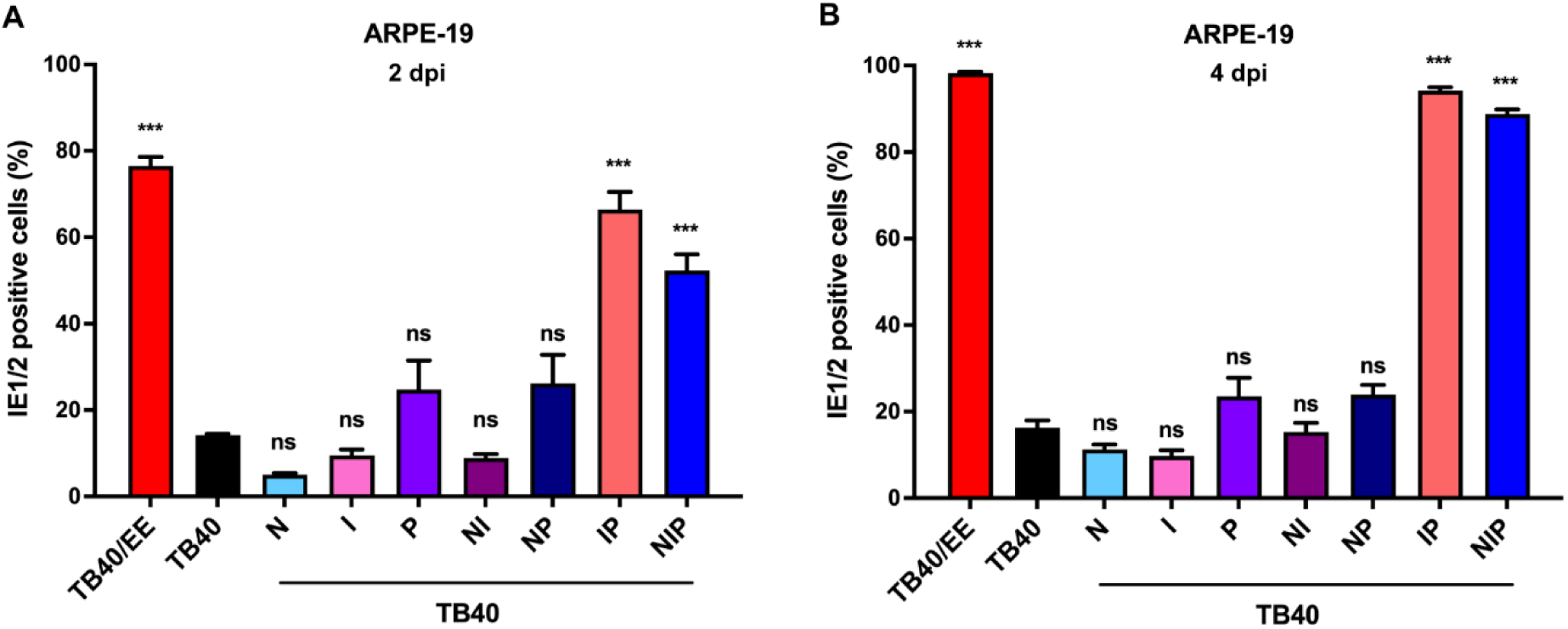
Relative infectivity of TB40/EE and recombinant TB40 strains. (A, B) ARPE-19 cells were infected at an MOI of 1 (based on titers determined on HFF). The relative infectivity of recombinant TB40 strains was assessed at 2 and 4 dpi by quantifying the percentage of IE1/2-positive nuclei. Mean ± SEM of three independent experiments. Recombinant TB40 strains were compared to TB40. Significance was determined by one-way ANOVA with Dunnett’s multiple comparison test. ***, *P* < 0.001; ns, not significant.

### An intronic SNV in UL128 increases transcript splicing but decreases infectivity and replication in epithelial cells

To assess the ability of the recombinant viruses to replicate and spread in ARPE-19 epithelial cells, ARPE-19 cells were infected at an MOI of 0.1 and virus released into the supernatant was measured by titration on ARPE-19 cells. Multistep replication kinetics (Fig. 4A and B) showed that TB40/EE replicated to slightly higher titers than TB40. Unexpectedly, TB40-P replicated to substantially lower titers than TB40 and the other derivatives (Fig. 4A). However, when the UL128 SNV was combined with the UL122 SNV (in mutants TB40-IP and TB40-NIP), viral replication was very similar to TB40/EE (Fig. 4B). To determine whether differences in viral infectivity were responsible for the impaired replication of TB40-P, we repeated the replication kinetics experiment with TB40/EE, TB40, TB40-P, and TB40-IP. Virus release into the supernatant was measured by both titration and quantitative PCR (qPCR) (Fig. 4C and D), and the infectivity of the different viruses was calculated as infectious units (IU) per 1000 genomes (Fig. 4E). The results suggested that TB40-P virions were substantially less infective than those of the other strains.

**Figure 4.**
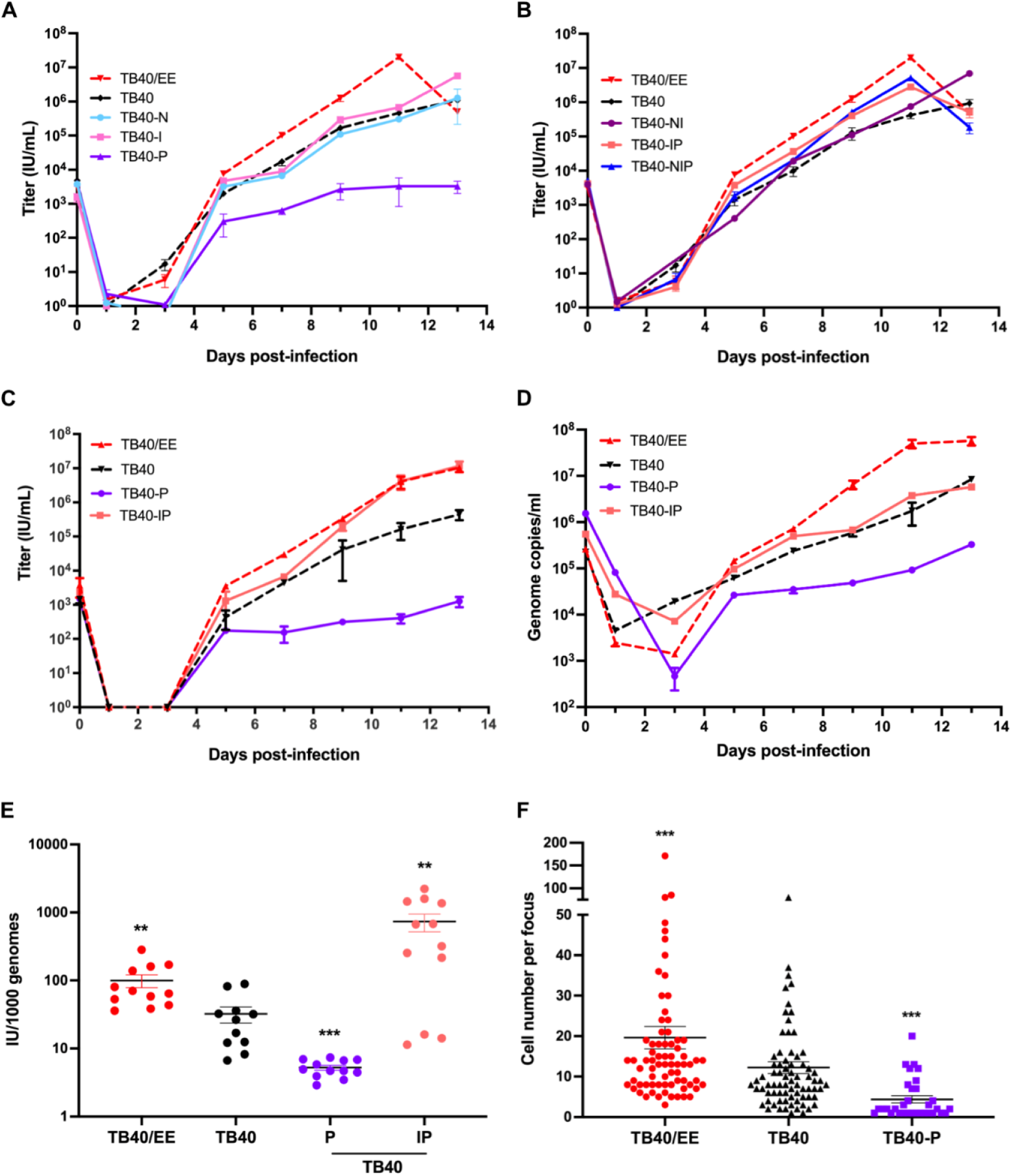
Replication and infectivity of recombinant TB40 strains in ARPE-19 cells. (A, B) ARPE-19 cells were infected at an MOI of 0.1. Supernatants were collected at various times post-infection and titrated on ARPE-19 cells. Titers are shown as the mean ± SEM of three biological replicates. (C-E) ARPE-19 cells were infected at an MOI of 0.05 with TB40/EE and recombinant TB40 strains. (C) Cell-free virus released into the supernatant was titrated to determine infectious units (IU). (D) Viral genome copies in the supernatant were quantified by qPCR. Mean ± SEM of three biological replicates are shown. (E) Infectivity was calculated as infectious units (IU) per 1000 viral genomes at days 7, 9, 11, and 13 post-infection with three biological replicates each. (F) ARPE-19 cells were infected at an MOI of 0.1. Cell-to-cell spread was evaluated using a focus expansion assay. At 5 dpi, cells were fixed and analyzed by immunofluorescence for IE1/2 expression. The number of infected cells per focus was determined by fluorescence microscopy and automated counting. Values were compared to TB40, and significance was determined using the Student’s *t*-test. **. *P* < 0.01; ***, *P* < 0.001.

We also compared TB40/EE, TB40, and TB40-P in a focus expansion assay to assess differences in cell-to-cell spread. ARPE-19 cells were infected at a low MOI, and focus size was determined at 5 dpi by counting IE1/2-positive nuclei per focus (Fig. 4F). Compared to TB40, average focus size was significantly higher for TB40/EE and significantly reduced for TB40-P, suggesting that differences in cell-to-cell spread contributed to the differences in replication observed in the previous experiments. To exclude the possibility that an adventitious mutation caused the reduced replication and spread of TB40-P, we isolated viral DNA from the TB40-P stock and sequenced the viral genome. No unintended mutations were detected.

The UL128 SNV is located within intron 1, close to the splice acceptor site (Fig. 5A). The SNV present in TB40-BAC4 was previously shown to reduce the splicing of UL128 exons 1 and 2 when introduced into the Merlin strain (32). To verify that this SNV also affects the splicing of UL128 in the TB40-BAC4 background, we determined the percentage of spliced transcripts at 3 and 5 dpi by qRT-PCR. As shown in Figures 5B and C, UL128 splicing was less efficient in TB40-infected cells than in those infected with TB40/EE, TB40-P, or TB40-IP. However, a comparison of UL128 protein levels in purified viral particles revealed that TB40/EE and TB40-IP virions contained more UL128 protein than TB40 and TB40-P virions (Fig. 5D). These data suggested that both UL128 and UL122 (encoding IE2), need to be modified in TB40-BAC4 to reproduce the properties of TB40/EE.

**Figure 5.**
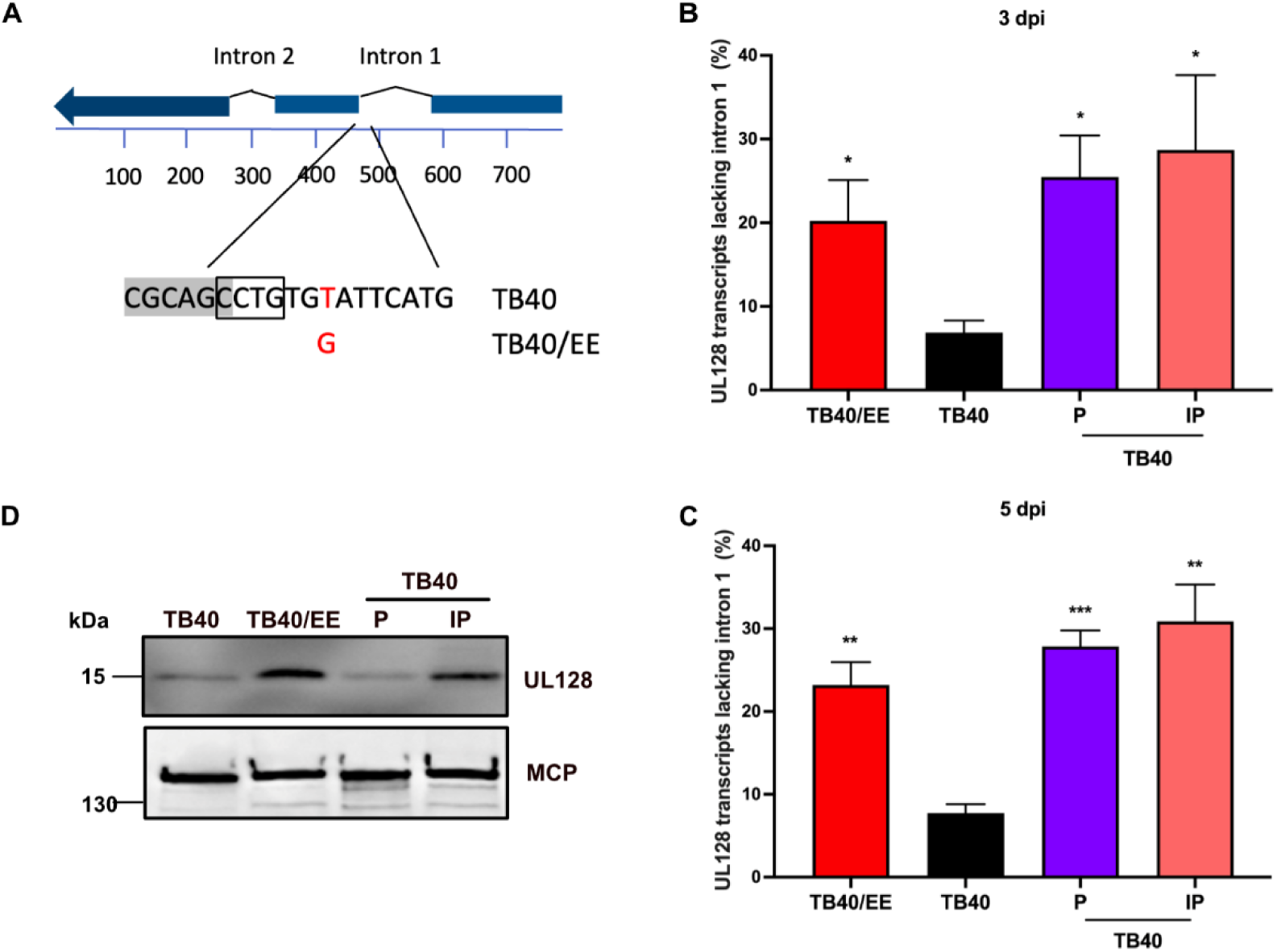
UL128 splicing is affected by an SNV in intron 1. (A) Schematic representation of the UL128 transcript and the SNV in intron 1. The consensus splice acceptor sequence is boxed. (B, C) ARPE-19 cells were infected with the indicated virus, and total RNA was extracted at 3 (B) or 5 days post-infection (C). Total UL128 transcripts and transcripts lacking intron 1 were quantified by qRT-PCR. The percentage of spliced *UL128* transcripts is shown as mean ± SEM of three biological replicates. (D) Purified viral particles were lysed and analyzed by immunoblot using antibodies against UL128 and the major capsid protein (MCP, loading control).

### The IE2 D390 variant affects early and late protein levels

IE2 is the main HCMV transactivator and regulates viral early and late gene expression (24). A previous study showed that the IE2 D390 variant in TB40-BAC4 allows this virus to replicate in the absence of UL84 (30). However, the underlying mechanism remains to be determined. To investigate whether the amino acid (D versus H) at position 390 affects the IE2-dependent expression of viral early and late genes, ARPE-19 cells were infected at an MOI of 2, using virus stocks titrated on the ARPE-19 cells to ensure equivalent infection. The accumulation of early and late proteins involved in viral replication (UL44, UL57, and UL84) or infection (gO and UL128) was analyzed at 1, 2, and 3 dpi. Strikingly, all viruses expressing the IE2 H390 variant (i.e., TB40/EE, TB40-I, and TB40-IP) expressed higher levels of UL44, UL57, and UL84 proteins than those expressing the D390 variant (TB40 and TB40-P) (Fig. 6). Interestingly, substantial amounts of UL128 protein were only detected in cells infected with TB40/EE or TB40-IP. These results suggested that the IE2 H390 increases the expression of several early and late proteins. However, increased UL128 protein levels require both the IE2 H390 variant and the SNV affecting *UL128* splicing.

**Figure 6.**
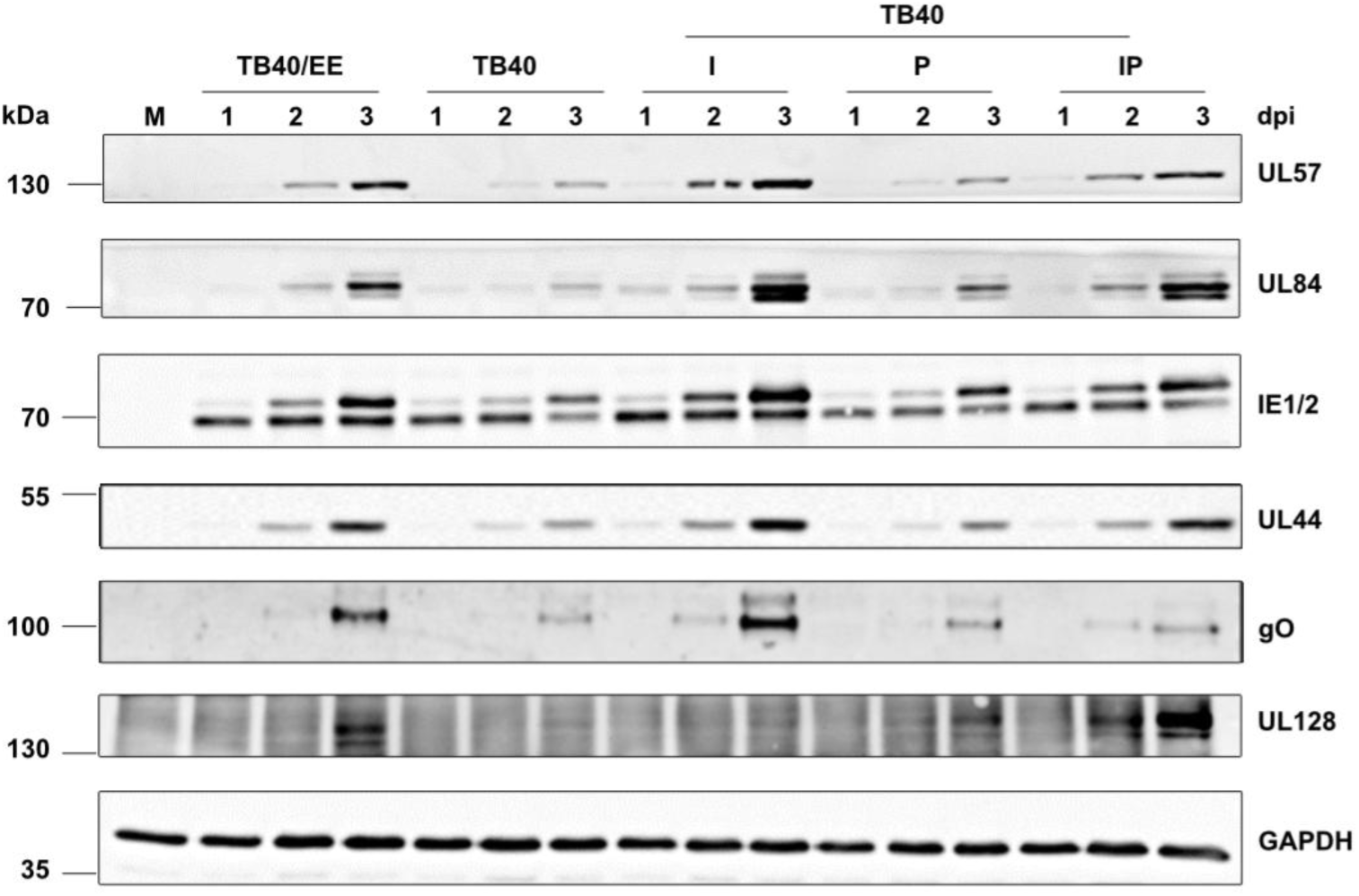
Accumulation of early and late proteins upon infection with TB40/EE and recombinant TB40 strains. ARPE-19 cells were infected at an MOI of 2 (based on titers determined on ARPE-19 cells). Cell lysates were collected at 1, 2, and 3 days post-infection (dpi). Expression levels of viral proteins involved in infection (gO, UL128) and replication (IE1/2, UL44, UL57, UL84) were analyzed by immunoblotting. A non-denaturing gel was used for the detection of UL128. All other proteins were separated on denaturing gels.

### SNVs in UL128 and UL122 jointly increase infectivity in macrophages and trophoblast cells

The previous experiments demonstrated that a combination of SNVs in UL128 and UL122 are responsible for the high infectivity of TB40/EE on ARPE-19 epithelial cells (Fig. 3). To test whether the same SNVs might also affect viral infectivity on other cell types, we determined the relative infectivity of the recombinant viruses on THP-1-derived macrophages and JEG-3 trophoblast cells. THP-1-derived macrophages were infected at an MOI of 0.5, while JEG-3 cells were infected at an MOI of 5. The percentage of IE1/2-positive cells was determined at 2 dpi by immunocytochemistry (Fig. 7A and B). As previously observed in ARPE-19 cells (Fig. 3), TB40-IP and TB40-NIP mutants had the highest infectivity, comparable to that of TB40/EE, suggesting that the same SNVs increase infectivity in different non-fibroblast cell types.

**Figure 7.**
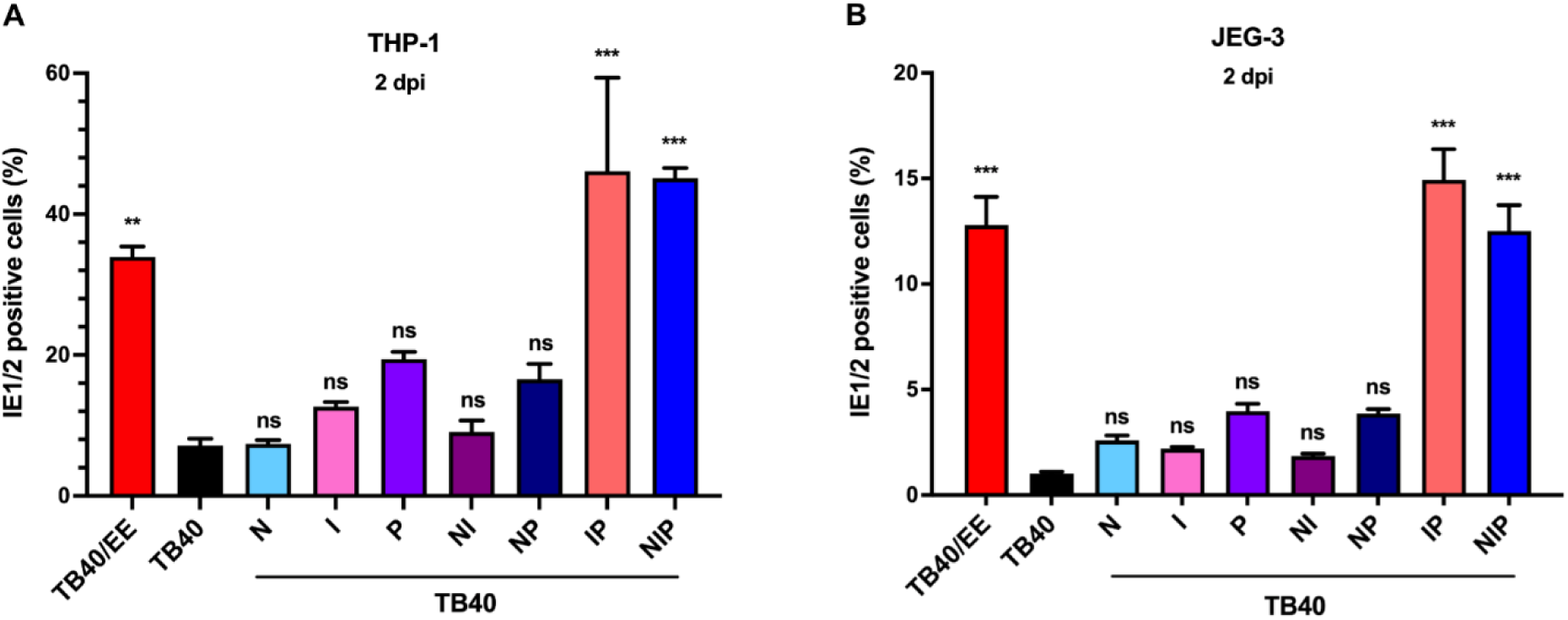
Relative infectivity of TB40/EE and recombinant TB40 strains on THP-1 and JEG-3 cells. (A) THP-1-derived macrophages were infected at an MOI of 0.5, and (B) JEG-3 cells were infected at an MOI of 5 (based on titers determined on HFF). The relative infectivity was assessed at 2 dpi by quantifying the percentage of IE1/2-positive nuclei. Mean ± SEM of three independent experiments are shown. Recombinant TB40 strains were compared to TB40. Significance was determined by one-way ANOVA with Dunnett’s multiple comparison test. **, *P* < 0.01; ***, *P* < 0.001; ns, not significant.

## Discussion

HCMV substrain TB40/EE was derived from the parental TB40/E strains by 3 sequential passages on ARPE-19 retinal epithelial cells and showed increased replication and syncytium formation (45). To identify the genetic determinants of these remarkable biological properties, we compared its sequence to that of TB40-BAC4. We identified a limited number of sequence variations and focused on three SNVs in UL26, UL122, and UL128. By introducing these TB40/EE-specific SNVs into TB40-BAC4 by BAC mutagenesis, we demonstrated that the SNV in *UL122* combined with the SNV in *UL128* are necessary and sufficient to confer high infectivity and the ability to induce syncytium formation to TB40. By contrast, the SNV in *UL26* (resulting in an E98K amino acid substitution) did not contribute to this phenotype.

The UL26 ORF encodes a viral tegument protein important for high-titer replication (50). It restricts the cytokine signaling pathways during infection and antagonizes NF-κB activation (49, 51) by interacting with the protein inhibitor of activated STAT1 (PIAS1), a crucial regulator of STAT and NF-κB transcription factors (52). Although the UL26 SNV was not strongly enriched in TB40/EE (46), the function of UL26 as a modulator of host innate immune signaling made it a strong candidate as a contributor to the TB40/EE phenotype. However, the introduction of the UL26 E92K substitution into TB40-BAC4 did not increase viral infectivity in different cell types (Figs. 3 and 7), nor did it promote virus-induced cell-cell fusion (Fig. 2).

A previous study identified a TB40-BAC4-specific SNV in intron 1 of UL128, which affects the splicing efficiency of the UL128 transcript (32). The authors correlated this SNV to the low UL128 protein levels present in TB40 virions and demonstrated that its introduction into the HCMV strain Merlin reduced UL128 protein levels in virions (32). However, as we show here, the reverse experiment yielded an unexpected result: change of the single-nucleotide in TB40-BAC4 was not sufficient to cause a substantial increase in UL128 protein levels in virus-infected cells and viral particles (Figs. 5 and 6). Even though TB40-P expressed slightly increased levels of UL128 in ARPE-19 cells (Fig. 6), it replicated and spread less efficiently than the parental TB40 in these cells (Fig. 4). How can this unexpected finding be explained? A recent publication showed that the pentameric complex inhibits HCMV IE gene transcription, possibly by activating receptor-dependent signaling (53). Thus, it seems likely that the improved splicing of the UL128 transcript dampens viral gene expression. However, this negative effect was reversed when the IE2-encoding UL122 ORF was modified as well. In cells infected with TB40-IP, the expression of several viral proteins (including UL128) was increased (Fig. 6) and UL128 protein incorporation into viral particles and was elevated (Fig. 5D). This led to an increased infectivity for epithelial cell, macrophages, and trophoblast cells (Figs. 3 and 7).

TB40-BAC4 encodes an IE2 D390 variant, whereas TB40/EE and most other strains (including AD169, Towne, and Merlin) encode a histidine at amino acid position 390. Remarkably, the D390 variant allows viral replication in the absence of UL84 (30), a protein previously thought to be essential for HCMV replication. UL84 and IE2 interact with each other and cooperatively activate the *ori*Lyt promoter, which drives UL57 expression (26). On the other hand, UL84 was shown to inhibit IE2-mediated promoter activation, suggesting that the IE2-UL84 interaction may also dampen viral gene expression (54). In the present study, we demonstrated that several viral early and late proteins are expressed at lower levels if the virus expresses the IE2 D390 variant (Fig. 6), suggesting that this variant is a weaker transactivator than the H390 variant. Alternatively, the amino acid at position 390 might alter the interaction with the UL84 protein. Additional studies, which extend beyond the scope of the present article, will be necessary to elucidate the precise roles of IE2 and UL84 in the cooperative regulation of viral gene expression and *ori*Lyt-dependent replication.

Is the IE2 D390 variant a quirk of nature or an adventitious mutation that occurred during the passage of TB40/E in cell culture? Besides TB40-BAC4, only one other HCMV isolate (P10, GenBank QPZ45048) encodes an IE2 D390 variant. However, similar to TB40-BAC4, the TR strain also replicates in a UL84-independent fashion (30). It encodes another IE2 variant (D547), which is also present in two clinical HCMV isolates (JER851, GenBank AKI23906; JER1289, GenBank AKI24414). Hence, IE2 variants that allow UL84-independent replication are likely to be naturally occurring variants rather than cell culture-induced mutations, and they may be more frequent than anticipated.

The results of this study show that the infectivity of TB40/E-derived strains correlates with the expression levels of UL128, a component of the pentameric glycoprotein complex. TB40/EE and TB40-IP express high levels of UL128 protein and are highly infectious for epithelial cells and macrophages. This is not surprising as the infection of these cell types is known to be pentamer-dependent. We further showed that the infection of JEG-3 trophoblast cells also dependents on pentamer levels (Fig. 7B). Previous studies reported different levels of infectability of human trophoblast cells with different HCMV strains (55, 56), suggesting that differences in pentamer levels are (at least in part) responsible for the observed differences.

Besides the trimeric and the pentameric complexes, a third glycoprotein complex has recently been identified (57). It consists of gH and the glycoproteins encoded by UL116 and UL141 and was named gH-associated tropic entry (GATE) complex. It enhances infection of endothelial cells and possibly other cell types (57). Interestingly, some HCMV strains have acquired inactivating mutations in UL141 during propagation in cell culture (31, 33), suggesting that the GATE complex might be detrimental to viral replication in certain cell types, just like the pentameric complex is detrimental to HCMV replication in fibroblasts. As TB40/EE and TB40-BAC4 both carry the same frame-shift mutation in UL141, this mutation was not considered as a cause of the different properties of the two viruses. However, another recent study reported that UL141 also interacts with the trimeric and pentameric complexes (58). Thus, it should be worthwhile to investigate in future studies how UL141 modulates viral infectivity for different cell types and how it affects infection-induced cell-cell fusion.

## Materials and Methods

### Cell Culture

Human foreskin fibroblasts (HFF-1, SCRC-1041), human embryonic lung fibroblasts (MRC-5, CCL-171), and retinal pigmented epithelial cells (ARPE-19, CRL-2302) were obtained from the American Type Culture Collection (ATCC). HFF-1 cells were cultured in Dulbecco’s Modified Eagle’s Medium (DMEM, PAN-Biotech), supplemented with 5% fetal calf serum (FCS, Gibco), 100 U/mL penicillin, and 100 μg/mL streptomycin (pen/strep). ARPE-19 cells were maintained in DMEM/F-12 GlutaMAX (Gibco) with 10% FCS, pen/strep, 15 mM HEPES, and 0.5 mM sodium pyruvate. For infection experiments, both cell types were cultured in DMEM medium containing only FCS and antibiotics. The THP-1 monocytic cell line (ACC 16) was purchased from the German Collection of Microorganisms and Cell Cultures (DSMZ). THP-1 cells were propagated in RPMI 1640 GlutaMAX (Gibco) supplemented with 10% FCS, pen/strep, 10 mM HEPES, 1 mM sodium pyruvate, and 50 μM β-mercaptoethanol. To induce differentiation into macrophages for infection experiments, THP-1 cells were treated with 50 nM phorbol 12-myristate 13-acetate (PMA, Sigma-Aldrich) for 24 hours. JEG-3 trophoblast cells (ATCC HTB-36) were kindly provided by Dr. Pietro Scaturro (Leibniz Institute of Virology, Hamburg, Germany). These cells were grown in RPMI 1640 GlutaMAX, supplemented with 10% FCS and pen/strep. After four hours of infection, the culture medium was replaced with RPMI 1640 GlutaMAX containing 2% FCS, pen/strep, and 1 ng/mL epidermal growth factor (Gibco) to support cell maintenance.

### Viruses and stock preparation

The HCMV strain TB40/EE was described previously (45, 46). Infectious virus was reconstituted from TB40-BAC4 (43) and derivatives by transfection of fibroblasts as described (59). To prepare cell-free virus stocks, ARPE-19 cells were infected at low MOI. Supernatants were collected from ARPE-19 cells exhibiting 100% cytopathic effect. Cellular debris was removed by centrifugation at 5,000 × *g* for 10 min. Viral particles were pelleted by ultracentrifugation at 23,000 rpm for 60 min at 4°C using a Beckman Coulter SW 28 rotor. Virus pellets were resuspended in DMEM containing a cryopreserving sucrose-phosphate buffer (74.62 g/L sucrose, 1.218 g/L K₂HPO₄, 0.52 g/L KH₂PO₄), aliquoted, and stored at -80°C.

Viral titers were determined as previously described (48). Briefly, HFF or ARPE-19 cells were inoculated with serial dilutions of the virus stocks. After 48 hours, cells were fixed with methanol/acetone for 10 minutes at -20°C, blocked with PBS containing 1% milk for 15 min at 37°C, and incubated with an anti-IE1/2 antibody for 2 hours at 37°C. This was followed by incubation with an HRP-conjugated rabbit anti-mouse secondary antibody (Jackson ImmunoResearch) for 45 minutes at 37°C. Staining was performed using 3-amino-9-ethylcarbazole (BioLegend) as a horse radish peroxidase substrate. IE1/2-positive nuclei were counted, and viral titers were expressed as infectious units per milliliter (IU/mL).

### Plasmids and Reagents

The plasmid pEPkan-S and *E. coli* strain GS1783 were previously described (60). GS1783 bacteria were cultured in Lennox LB broth (Carl Roth) supplemented with 5 g/L NaCl (Sigma-Aldrich). Lentiviral vector plasmids encoding DSP1-7 or DSP8-11 and helper plasmids pMD-G and pCMVR8.91 were provided by Dalan Bailey (The Pirbright Institute, Woking, UK). Antibiotics, including ampicillin (100 µg/mL), kanamycin (50 µg/mL), and chloramphenicol (15 µg/mL), were obtained from Roth or Invitrogen. L-(+)-arabinose was purchased from Sigma-Aldrich.

### Mutagenesis of HCMV genomes

Point mutations in specific regions of interest were introduced into TB40-BAC4 by *en passant* mutagenesis (60). All modified BACs were validated by restriction fragment analysis and sequencing of the mutated region. To further verify integrity and absence of unintended mutations, the complete genomes of TB40-P and TB40-IP were sequenced using Illumina technology at the Next Generation Sequencing facility of the Leibniz Institute of Virology, following established protocols (61). The complete genome sequences of TB40/EE (MW439039) and TB40-BAC4 (EF999921) are available in GenBank.

### Replication kinetics

ARPE-19 cells were seeded one day prior to infection in 6-well plates at a density of 2 × 10⁵ cells per well. Cells were infected at an MOI of 0.1 for 3 hours at 37°C, followed by washing with PBS (Sigma-Aldrich) and the addition of fresh medium. Supernatants were collected at different times post-infection and titrated as described above.

### Antibodies

The following monoclonal antibodies were used in this study: anti-IE1/2 (3H4) was provided by Thomas Shenk (Princeton University, USA), anti-MCP (28–4) by William Britt (University of Alabama, USA), and anti-UL128 (4B10) by Barbara Adler (University of Munich, Germany). Anti-gO (clone gO.01) was purchased from the Center for Proteomics (Rijeka, Croatia), anti-UL57 and anti-UL84 from Santa Cruz, anti-UL44 (10D8) from Virusys, and anti-GAPDH (14C10) from Cell Signaling.

### Immunoblot detection

Immunoblotting was performed following standard protocols. ARPE-19 cells were seeded in 6-well plates (2 × 10^5^ cells/well) and infected on the following day at an MOI of 2. Cells were harvested at the indicated time points and lysed with RIPA buffer (50 mM Tris-HCl, pH 8, 150 mM NaCl, 1 mM EDTA, 1% NP-40, 0.5% deoxycholate, 0.1% SDS) supplemented with cOmplete Mini Protease Inhibitor Cocktail (Roche). For reducing conditions, 10% β-mercaptoethanol was added to the loading buffer. Equal protein amounts of each sample were separated by SDS-PAGE, transferred onto nitrocellulose membranes (Amersham) by semidry blotting, and detected using specific primary antibodies followed by HRP-conjugated secondary antibodies (Jackson ImmunoResearch) and enhanced chemiluminescence substrate (Amersham) with the addition of 10% Lumigen TMA-6 (Bioquote Limited).

For protein detection in virions, cell-free virus stocks were further cleared by centrifugation at 1000 × *g* for 5 min at 4°C and resuspended in RIPA buffer containing protease inhibitors. Insoluble material was cleared by an additional centrifugation at 16,000 × *g* for 10 min, and the resulting extracts were boiled at 95°C for 5 min in loading buffer.

### Immunofluorescence

For immunofluorescence analysis, ARPE-19 cells were seeded onto 8-well μ-slides (Ibidi) and infected on the next day. At 3 dpi, cells were fixed with methanol/acetone, blocked with PBS/1% milk, and incubated with primary antibodies either overnight at 4°C or for 2 hours at 37°C. Detection was carried out using an AlexaFluor 555-conjugated secondary antibody (Thermo Fisher Scientific). Nuclear DNA was counterstained with DAPI (4′,6-diamidino-2-phenylindole, Sigma-Aldrich). Imaging was performed using a Nikon A1+ LSM confocal microscope, and infectivity assays were conducted on a CellInsight CX5 High-Content Screening Platform (Thermo Fisher Scientific). The percentage of IE1/2-positive cells was then quantified using HCS Studio software.

### Quantification of HCMV genomes

Cell-free HCMV DNA was extracted from 200 µL supernatant using the InnuPREP DNA Mini Kit (Analytik Jena) and eluted in 50 µL nuclease-free water. Viral genome copies were quantified by real-time quantitative PCR (qPCR) using SYBR Green dye (Thermo Fisher Scientific) and primers specific for the HCMV UL36 ORF (ACGCAAAGAGTTCCTCGTAC and TGAACATAACCACGTCCTCG). PCR amplification was performed using a QuantStudio 3 qPCR machine (Thermo Fisher Scientific). For each experiment, two independent viral DNA extractions and three independent qPCR reactions were conducted.

### Quantitative real-time RT-PCR analysis

To quantify UL128 transcripts, ARPE-19 cells were infected at an MOI of 3. Total RNA was extracted from infected cells at 3 and 5 days post-infection using the InnuPREP RNA Mini Kit (Analytik Jena). Contaminating DNA was removed using the TURBO DNA-free Kit (Ambion). cDNA was synthesized from 1 µg of extracted RNA using RevertAid H Minus Reverse Transcriptase, oligo-dT primers, and the RNase inhibitor RiboLock (Thermo Fisher Scientific). qPCR was performed using the QuantStudio 3 Real-Time PCR System (Thermo Fisher Scientific) with 10 ng of cDNA and PowerTrack SYBR Green Mastermix (Thermo Fisher Scientific). UL128 transcript levels were normalized to IE1 at 1 day post-infection and the housekeeping gene GAPDH.

Primer sequences used for IE1, UL128, and GAPDH are as follows: IE1: (CAAGTGACCGAGGATTGCAA and CACCATGTCCACTCGAACCTT), UL128 (lacking intron 1): (CATGGTGGTGACGATCCCGCG and ATCGCTTCACCGTCGCGCTG), UL128 (all isoforms): (GCGGGTGGTTGACGTTTATG and AACCTGACGCCGTTCTTGA), and GAPDH: (CCCACTCCTCCACCTTTGACG and GTCCACCACCCTGTTGCTGTAG).

### Cell-cell fusion assay

ARPE-19 cells expressing the dual split protein (DSP) system were generated by transduction with lentiviral vectors, following a method described previously (47). Briefly, HEK293T cells were transfected with the lentiviral vector plasmid and helper plasmids pCMVR8.91 and pMD-G. Lentivirus-containing supernatants were then used to transduce ARPE-19 cells, and the transduced cells were selected using 1.5 µg/mL puromycin (Sigma-Aldrich).

For the fusion assay, equal numbers of ARPE-19 cells expressing DSP1-7 or DSP8-11 were combined and infected with HCMV at an MOI of 1. At 5 dpi, cells were washed with PBS and incubated with coelenterazine-h (Promega) at a final concentration of 2.5 nM. *Renilla* luciferase activity was quantified by measuring luminescence using a multi-mode microplate reader (FLUOstar Omega, BMG Labtech).

### Infectivity Assay

One day prior to infection, ARPE-19 cells and JEG-3 cells were seeded in 96-well plates at a density of 1 × 10⁴ cells/well, while macrophages were seeded at a density of 5 × 10⁴ cells/well. ARPE-19 cells were infected with HCMV at an MOI of 1, macrophages at an MOI of 0.5, and JEG-3 cells at an MOI of 5. The inoculum was removed at 3 hours post-infection, and cells were washed with PBS and incubated with fresh medium. At 2 and 4 dpi, infected cells were fixed and stained with an anti-IE1/2 antibody and DAPI. Fluorescence images were acquired, and the percentage of IE1/2-positive nuclei was determined with a CellInsight CX5 High-Content Screening Platform.

### Statistical Analysis

Statistical analyses were performed using GraphPad Prism 5.0 Software. Significance was determined using the Student’s *t*-test or one-way ANOVA followed by Dunnett’s multiple comparison test.

## Acknowledgments

We thank Thomas Shenk, William Britt, and Barbara Adler for antibodies, Dalan Baley for lentiviral vector plasmids, Jan Knickmann and Sanamjeet Virdi (LIV Next Generation Sequencing), as well as Roland Thünauer and Marcel Schie (LIV Microscopy and Image Analysis) for their help and support. Schematic figures were created with BioRender.

This research was supported by the Deutsche Forschungsgemeinschaft (grant BR1730/9-1 to W.B.). X.Z. and L.Y. were supported by scholarships from the China Scholarship Council (No. 202008080174 and 202106910030) and G.C. by a Leibniz Center Infection (LCI) scholarship.

Conceptualization, L.H. and W.B.; methodology, X.Z., G.C., and G.F.; investigation, X.Z., L.Y., G.C., and A.A.-P.; writing—original draft preparation, X.Z. and W.B.; writing—review and editing, all authors; supervision, G.F. and W.B.; funding acquisition, G.F. and W.B.

## Data availability

The complete sequences of HCMV strain TB40-BAC4 and TB40/EE are available at GenBank (EF999921 and MW439039). All other data supporting the findings of this study are contained within the article.

